# Exploiting a Mechanical Perturbation of Titin Domain to Identify How Force Field Parameterization Affects Protein Refolding Pathways

**DOI:** 10.1101/764076

**Authors:** David Wang, Piotr E. Marszalek

**Affiliations:** Department of Mechanical Engineering and Materials Science, Duke University, Durham, NC 27708, USA

## Abstract

Molecular mechanics force fields have been shown to differ in their predictions of processes such as protein folding. To test how force field differences affect predicted protein behavior, we created a mechanically perturbed model of the beta-stranded I91 titin domain based on atomic force spectroscopy data and examined its refolding behavior using six different force fields. To examine the transferability of the force field discrepancies identified by this model, we compared the results to equilibrium simulations of the weakly helical peptide Ac-(AAQAA)_3_-NH_2_. The total simulation time was 80 *µs*. From these simulations we found significant differences in I91 perturbation refolding ability between force fields. Concurrently, Ac-(AAQAA)_3_-NH_2_ equilibration experiments indicated that although force fields have similar overall helical frequencies, they can differ in helical lifetimes. The combination of these results suggests that differences in force field parameterization may allow a more direct transition between the beta and alpha regions of the Ramachandran plot thereby affecting both beta-strand refolding ability and helical lifetimes. Furthermore, the combination of results suggests that using mechanically perturbed models can provide a controlled method to gain more insight into how force fields affect protein behavior.

## Introduction

Molecular dynamics simulations describe macromolecular motion in all-atom detail, a feat that is extremely difficult to achieve experimentally. ^1–5^ From these simulations, a wide variety of thermodynamic and kinetic observables can be inferred thereby providing a better picture of the nature of various biological processes such as protein folding.^6–12^ The utility of molecular dynamics simulations however, depends on the accuracy of its force field - the set of potentials used to describe the energetics behind atomic movements. Due to the complexity of the parameterization process, various force fields are not guaranteed to exhibit the same effect on biomolecular behavior.^13–26^ This has been seen previously when modifications to torsion angle parameters have been shown to impede folding and generate incorrect structures. ^27–29^ Thus force field variations can affect sensitive biomolecular processes such as protein folding pathways.^2,7,12,15,30,31^

Although current validation schemes of molecular mechanics force fields use a wide variety of experimental data to ensure transferability,^32–37^ these validations often occur over small peptides. Other studies involving the folding of complete proteins are often limited by simulation time scales^19,20,33^ or involve the use of enhanced sampling techniques like replica exchange.^2,7,9,32^ Thus, examining more detailed pictures of processes like protein folding that could also be simulated in a small time would be helpful in identifying force field discrepancies as these processes are more sensitive to force field inaccuracies than validation and comparison studies on native state structures. ^9,12,30^

One potential way to experimentally observe the behavior of macromolecular structures is through atomic force spectroscopy. ^38–40^ With atomic force microscopes, various proteins can be unfolded using force. Indeed, certain portions of proteins have been observed to unfold under force and refold after force relaxation.^38,41,42^ Thus, a force perturbation on a small portion of protein could serve as an ideal model to examine how various force fields comparatively affect biomolecular motion. Since the perturbations refold after force relaxation, we expect that a reconstruction of the structural perturbation will refold if the molecular mechanics force field can accurately reproduce the perturbation.

In particular, the structural perturbations involved in many atomic force microscopy refolding experiments are relatively tiny compared to the overall structure. ^38,42^ Therefore, by the thermodynamic hypothesis it is expected that structural perturbations will robustly refold into the native structure. ^5,29,43^ Furthermore the small size of the perturbation allows for faster computation because of the reduction of the size of the water box. Along with an increase in speed of computation, the perturbations should also be more sensitive to force field discrepancies as the perturbations should refold from the unfolded state to the native structure. ^9,12,30,44^

From these considerations, we created a model of a mechanically perturbed protein from titin I91 (formerly I27), a domain of the titin protein that has been extensively studied using force spectroscopy.^26,45–49^ Under force, the first ten residues of I91 comprising beta strand A have been shown to unfold and refold upon force extension. ^38^ Thus, I91 provides an opportunity to test the refolding of beta proteins with various force fields. In particular, this mechanical perturbation provides a more controlled method to examine the refolding of a single beta strand as compared to the refolding of beta-hairpins. Previous literature has indicated that beta-hairpin refolding occurs under multiple stages and the particulars of the folding process are still contested. ^13,19^ Furthermore, beta-strand refolding may require timescales greater than the simulation timescale of 1 *µs*.^7,9,13,19^ Thus, by examining the movement of a single strand, this mechanical perturbation provides a more controlled and faster method to examine the dynamics of beta strands.

The Ac-(AAQAA)_3_-NH_2_ helical peptide was chosen to model the folding and unfolding of helix structures. This helical peptide has been used as a benchmark to determine empirical corrections to various force fields^30,32,34^ and previous NMR data also provides an empirical observable with which helical frequencies can be validated. ^50^ The model provides a way to test whether force field discrepancies observed with the I91 perturbation are transferable and whether such observations can provide new insight into the folding process.

We tested five different force field/water model combinations using Amber simulation software: Amber ff14SB/Tip3p, ^33^ Amber ff14SB without backbone dihedral correction/Tip3p (abbreviated as Amber ff14SBonlySC), ^33^ Amber FB15/Tip3p, ^36^ Amber ff99SB*-ILDN/Tip3p, ^32^ and Amber ff99SB-ILDN/Tip3p; ^1^ and two different force field/water model combinations using Desmond: Charmm 22*/Tip3p-Charmm^30^ and Amber ff99SB-ILDN/Tip3p ^1^ for time spans amounting to at least 1 *µs* per simulation. The 1 *µs* timescale allowed for the observation of folding events while being short enough to run repeated trials to ensure robust sampling of the unfolded state. From these simulations, we observed differences in the I91 perturbation refolding pathway which were then supported by Ac-(AAQAA)_3_NH_2_ equilibration simulations. This suggests that mechanical perturbations may serve as good models to examine the effect of force field discrepancies.

## Simulation Methodology

In order to generate the structure perturbation, steered molecular dynamics simulations were conducted with NAMD. ^51^ The N-terminus of the I91 domain (using chain A of pdb structure 1waa, residues 1-89^45^) was pulled in the upwards z-direction away from the rest of the protein using a previously established protocol.^46^ A structure was selected when residues 1-10 had lost contact with the rest of the I91 domain. The upward z-direction was chosen in order to examine the behavior of the strand in the unfolded state as z-directional pulling leaves more space for the strand to sample various configurations. Although the direction of pulling is not the same as that done during experiment, we expect that given the small size of the perturbation and the downhill nature of the folding process^29,43^ that the perturbation will still exhibit robust refolding. The selection of these structures was then used as the starting point of equilibration simulations.

Equilibration simulations of the I91 perturbation were performed using two different software packages. A water box was constructed around the initial structure with a buffer of 10 Å on each side. In order to test the Charmm 22* and Amber ff99SB-ILDN force fields, the Desmond software package was used alongside GPU acceleration.^52^ Using Desmond, the initial perturbed structure was equilibrated at 310K with a Nose-Hoover thermostat in an NPT ensemble for an initial 1*µs* of simulation time. If the structure did not reach the folded state by the end of the simulation, the simulation was continued for a second 1*µs* of simulation time. Five trials were performed for both the Charmm 22* and Amber ff99SBILDN force field with each trial changing the randomization seed to provide independence. In order to test the Amber ff14SB, Amber ff14SBonlySC, Amber FB15, Amber ff99SB*ILDN, and Amber ff99SB-ILDN force fields the Amber simulation package was used alongside GPU acceleration^53^ Using Amber, the perturbed structure was equilibrated at 310K with a Langevin thermostat (friction of 2 ps^*−*1^) in an NPT ensemble for an initial 1 *µs* of simulation time. Similarly, if the structure did not reach the folded state by the end of the initial 1 *µs* of simulation time, the simulation would be continued for another 1 *µs*. Five trials were performed for each of the Amber ff14SB, Amber ff14SBonlySC, Amber ff99SB*-ILDN, Amber ff99SB-ILDN and Amber FB15 force fields.

Equilibration simulations of Ac-(AAQAA)_3_-NH_2_ were performed at 303K to test whether differences in force fields captured by the mechanical perturbation refolding simulations could also be observed in Ac-(AAQAA)_3_-NH_2_ helical distributions. For simulations using Amber software, the Ac-(AAQAA)_3_-NH_2_ structure was created as a linear peptide using tleap. The structure was placed in a water box with a 20 Å buffer and equilibrated using an NPT ensemble with a Langevin thermostat (friction of 1 ps^*−*1^) for 2*µs* for the following force fields: Amber ff14SB, ^33^ Amber ff14SBonlySC, ^33^ Amber ff99SB*-ILDN, ^32^ Amber FB15,^36^ and Amber 99SB-ILDN.^54^ For simulations using Desmond software, the Ac-(AAQAA)_3_-NH_2_ structure was again created as a linear peptide using tleap and run in an NPT ensemble with a Nose-Hoover thermostat for 2 *µs* for the following force fields: Charmm 22* ^30^ and Amber ff99SB-ILDN. ^1^ Unlike Amber software, Desmond software does not allow the automatic addition of NH_2_ caps therefore the c-terminus of Ac-(AAQAA)_3_-NH_2_ peptide was manually adjusted to fit a residue template in the Charmm 22* force field (see Supplemental Information), while in the Amber ff99SB-ILDN force field an NMA cap was used instead.

To determine how force fields affected the I91 perturbation refolding process, molecular dynamics trajectories were divided according to whether the structure was in the folded or unfolded states. To locate the folded and unfolded states, RMSD and Q-value plots for the 10 perturbed residues of the I91 perturbation were created with cutoffs of 0.4 and 0.85 being used for the folded and unfolded states respectively (see Supplementary Information). ^29^ Upon identification of the time of folding, the structures in the unfolded state were grouped from each of the five trials and analyzed to see whether force fields affected the unfolded state. Analysis algorithms were used to produce the potential of mean force distribution, ^55^ secondary structure percentage, ^56^ and clustered structures. ^57^ Secondary structure percentage was analyzed via DSSP^56^ using MDTraj.^58^ Clustering of the unfolded structures was done using the KMedoids and KCenters algorithms as implemented in MSMBuilder^59^ and MDTraj.^58^ Arrangement of clusters for visualization was made using the Multialign viewer ^60^ in UCSF Chimera. ^61^

To examine Ac-(AAQAA)_3_-NH_2_ helicity we used previously established criteria to define the a_*h*_ region: *φ ϵ*[−100°, −30°] and *ψ ϵ*[−67°, −7°] and counted a residue as helical if three consecutive residues were found in the a_*h*_ region.^32,34^ We computed the frequency with which each individual residue was deemed helical in order to match previously published results and NMR data.^30,32,34,50^ The helicity of each Ac-(AAQAA)_3_-NH_2_ residue as a function of time was computed using DSSP criteria^56^ with an implementation found in MDTraj. ^58^ The lifetime of each helix was defined as the length of time which a set of helical residues and their adjacent partners could propagate in the secondary structure vs time plot generated by DSSP (see Supplemental Information). These counts were then used to compute the mean helical lifetimes and max helical lifetimes of the simulation. The helical lifetime per residue was defined as the length of time with which a specific residue could remain helical (see Supplemental Information). From this set of times, the mean and max helical lifetimes per residue were also computed. Finally, dihedral potential of mean force plots were created in order to examine the distribution of *φ ψ* values found in the Ac-(AAQAA)_3_-NH_2_ equilibrium simulation distributions.^55^

## Results

### I91 refolds robustly under most force fields

From the simulations it can be seen that the perturbed structure of I91 domain refolds robustly under most force fields (see Table 1). For most force fields, the refolding times range from tens of nanoseconds to over 1 *µs*. This distribution suggests that while the given 2 *µs* of simulation time is feasible to capture some refolding events, the true range of refolding times is somewhat greater than the simulation time of 2 *µs* which was the maximum possible simulation time given the computational power at our disposal. There were noticeable differences in the ability of various force fields to capture refolding events. In particular, the Amber ff14SB and Amber ff99SB*-ILDN force fields displayed difficulty refolding the perturbed intermediate in the allotted time (see Table 1). This difference in refolding capacity suggests that differences in force field parameterization may affect the refolding pathways of I91. Since the unfolded state is more sensitive to force field discrepancies, we then examined the unfolded state to see if different force fields alter the unfolded state of the I91 mechanically perturbed intermediate.

**Table 1:** I91 Perturbation Refolding Simulation Summary.

| Force Field | Trials Refolded | Refolding Times (ns) |
| --- | --- | --- |
| Charmm 22* | 5/5 | 498, 749, 1089, 346, 1136 |
| Amber ff99SB-ILDN (Desmond) | 2/5 | 1372, 1270 |
| Amber ff99SB-ILDN (Amber) | 3/5 | 411, 833, 455 |
| Amber FB15 | 4/5 | 1384, 1208, 890, 57 |
| Amber ff14SB | 1/5 | 600 |
| Amber ff14SBonlySC | 3/5 | 852, 384, 51 |
| Amber ff99SB*-ILDN | 2/5 | 439, 1602 |

After separating the molecular dynamics trajectories into folded and unfolded states, we examined the distribution of dihedral angles in the unfolded state. Potentials of mean force were constructed in order to examine the magnitude with which the dihedral angles were found in various states (see Figure 2). From these potential of mean force diagrams it can be seen that there are significant minima in the helical regions of the Charmm 22*, Amber ff99SB-ILDN, Amber ff99SB*-ILDN, and Amber ff14SB force fields. The difference between the Amber ff14SB and Amber ff14SBonlySC force fields indicates that the empirical dihedral correction of Amber ff14SB force field sharply increases the amount of helical torsion angles found within the unfolded region by almost 2 *k*_*B*_T. Additionally, it is interesting that while the Charmm 22* force field appears to refold I91 robustly, the Amber ff99SB*-ILDN and Amber ff14SB force fields struggle to refold the perturbation. Since the Charmm 22* force field also contains a significant population of dihedral angles in the alpha helical region, this suggests that simply possessing residues with helical dihedral angles may not actually affect refolding.

**Figure 1:**
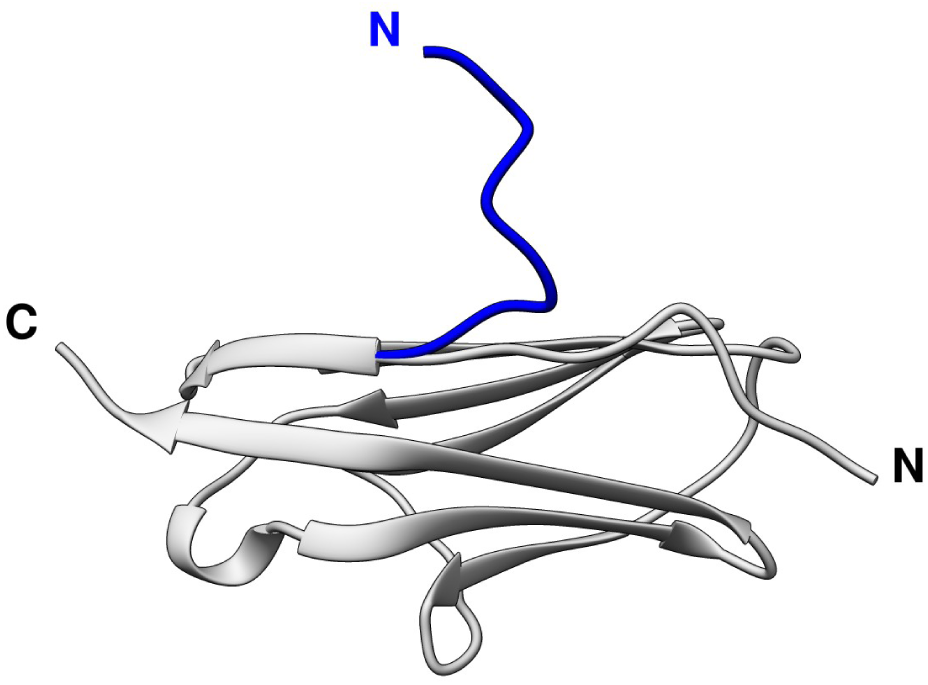
I91 Mechanical Perturbation. The native structure of I91 (PDB ID: 1waa ^45^) is shown in gray while the mechanical perturbation is shown in blue.

**Figure 2:**
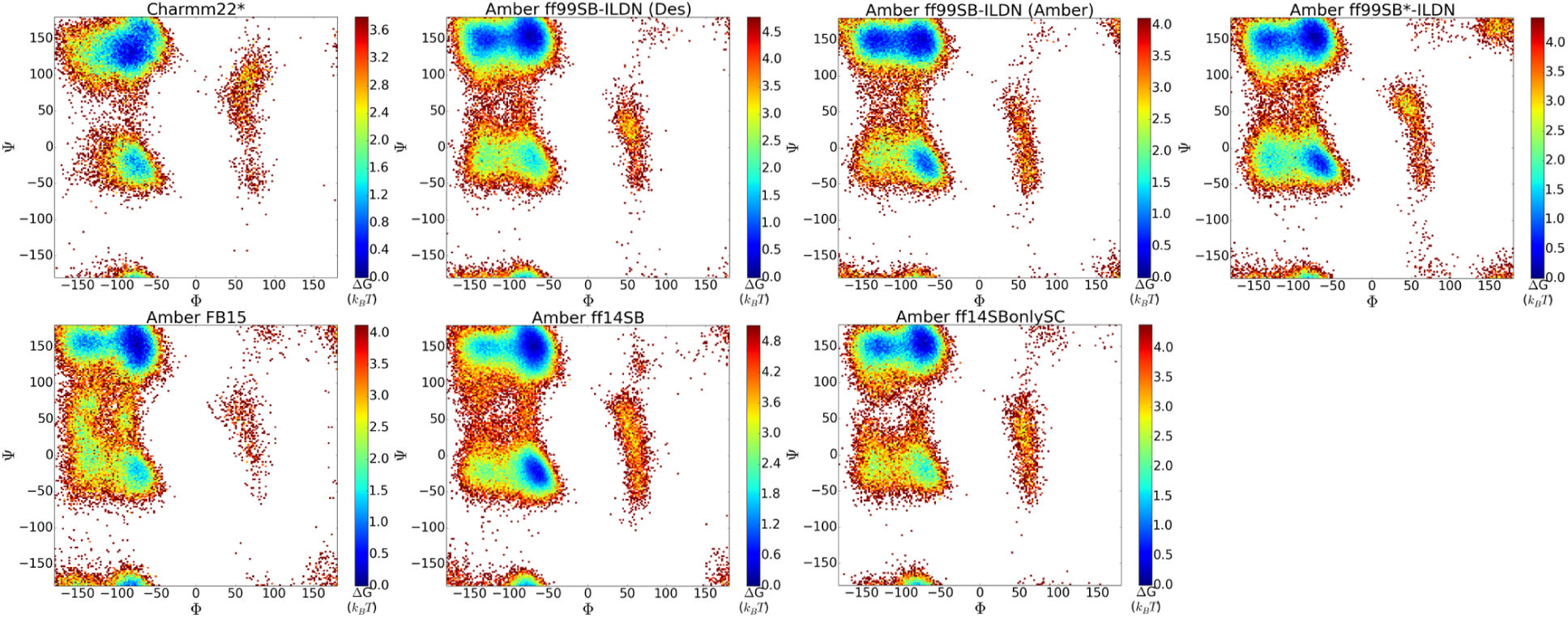
Potentials of mean force for the unfolded state of the I91 perturbation for each force field.

Another interesting point of comparison lies between the Amber FB15 force field and all the other force fields. Dihedral sampling with the Amber FB15 force field for the I91 unfolded state is much more prevalent in the region between the alpha and beta regions of the Ramachandran plot (*ψ ϵ* [25°,75°]). The potential of mean force plot for the unfolded state of the Amber FB15 force field then samples dihedral pairs which are not found in any of the other force fields (see Figure 2). This may be related to the parameterization method of the Amber FB15 force field which reparameterized all the bonded terms of the Amber ff99SB force field based on an automated methodology,^35^ notably relaxing bond force constants from their equilibrium values by up to an extra 10 percent. ^36^ The effects of this methodology are visible in the I91 potential of mean force map as the Amber FB15 force field samples more regions of the Ramachandran plot than any other force field.^13,36^

Finally, a point of comparison arises between the Amber ff99SB-ILDN force field simulations conducted with Desmond software and the Amber ff99SB-ILDN force field simulations conducted with Amber software. In comparison to the rest of the potential of mean force plots, the Amber ff99SB-ILDN potential of mean force plot is similar in shape to both the Amber ff14SB and Amber ff14SBonlySC potential of mean force plots. This is because the same backbone dihedrals are used for all three force fields - each being based on the Amber ff99SB backbone dihedrals.^1,33,62^ By comparing the two Amber ff99SB-ILDN force fields together it becomes apparent that the Amber ff99SB-ILDN force field simulations conducted with Amber possess a greater amount of helical torsion angles than the Amber ff99SB-ILDN force field using Desmond. However, given the higher amount of refolding events produced by the Amber ff99SB-ILDN force field simulations conducted with Amber, we believe that this is more likely to be the result of differences due to sampling than it is of any innate differences between the software packages.

Even though dihedral maps give a good idea of which residues possess helical or beta-sheet characteristics the actual formation of helices and beta-sheets does not occur until multiple residues have adopted the same repeating dihedral angles. Thus, in order to examine the actual secondary structure content of the unfolded state. We used DSSP to assign the secondary structure of each residue and then compiled the secondary structure frequencies together from all the unfolded states of all the trials (see Figure 3). By comparing the various secondary structures for the unfolded state of the I91 perturbation we can see that the Amber ff14SB and Amber ff99SB*-ILDN force fields possess a much higher frequency of helical residues than the rest of the force fields which produce almost no helical residues. This supports the potential of mean force plots (see Figure 2) which suggest large minima in the helical regions of both Amber ff14SB and Amber ff99SB*-ILDN. However, the absence of helices notably in Charmm 22* suggests that residues simulated using the Charmm 22* force field could easily leave the alpha helical minima region and did not remain in the helical configuration long enough for the formation of stable helices.

**Figure 3:**
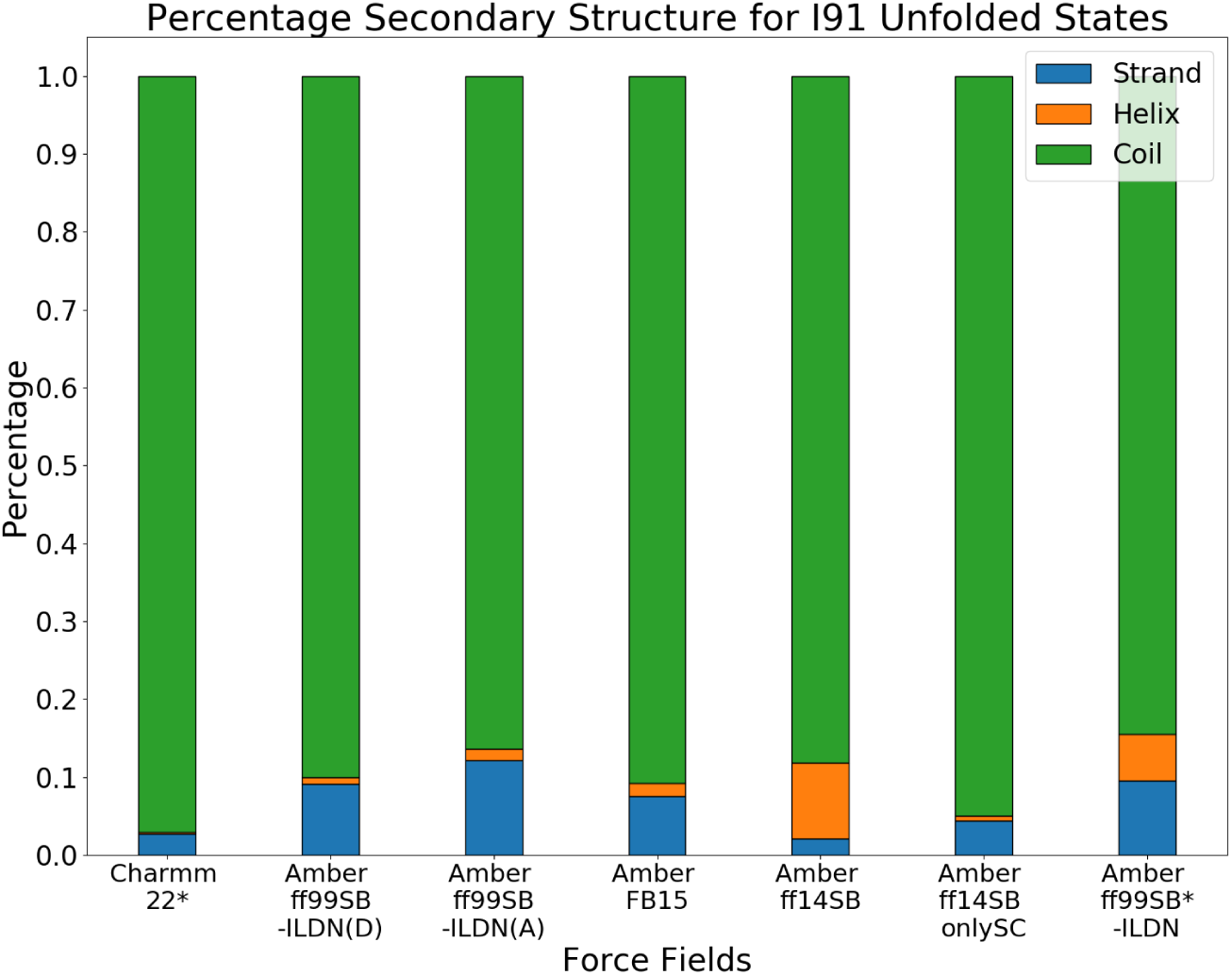
Secondary structure percentages for the unfolded states of the I91 perturbation. A reduced characterization is shown where a helix is either an alpha, 3/10, or pi helix; a strand is either a beta bridge or extended strand; and a coil is either a turn, bend, or irregular element.

Further comparisons can be made between the Amber FB15 force field and the rest of the structures. Overall, the Amber FB15 force field displays roughly similar amounts of secondary structure as the other force fields. This suggests that although the dihedral angle distributions are different, the FB15 force field will end up producing roughly equivalent amounts of secondary structure as the rest of the force fields. It is interesting to note that although most Amber force fields have a significant amount of extended conformation, that the Amber ff14SBonlySC and Charmm 22* force fields both possess low amounts of extended conformation. This suggests that the force fields do not form strands in the unfolded state and only adopt the secondary structure when folded into the native state.

After identifying common secondary structures, we wished to examine the most common structures sampled by each of the force fields. To that end we used the KMedoids and KCenters clustering algorithms to determine the most common structures of the unfolded state. Two different clustering algorithms were used because the KCenters clustering algorithm commonly picks up outliers in the structures and was thus used to identify the range of structures that were sampled in the unfolded state. ^57^ Meanwhile, the KMedoids algorithm was used to find the most common unfolded state structures and thus the structures that were most likely to be found in local minima.

Through this method we observed that the Amber ff14SB force field, while sampling much of the same space as the other force fields (see KCenters structures for Amber ff14SB in Figure 4e), possesses many structures found in helices as seen with the KMedoids algorithm (see Figure 4e). This corroborates the potential of mean force plots and secondary structure analysis using DSSP. The heavy amount of helical structures suggests that the sampling of this minima with the Amber ff14SB force field might temporarily impeded the creation of beta structures thereby altering the refolding pathway of I91. The presence of helical segments in the Amber ff99SB*-ILDN force field also supports the refolding times of this force field (see Table 1). The various helical residues of the Amber ff99SB*-ILDN force field may also impede refolding and cause the formation of unnecessary local minima.

**Figure 4:**
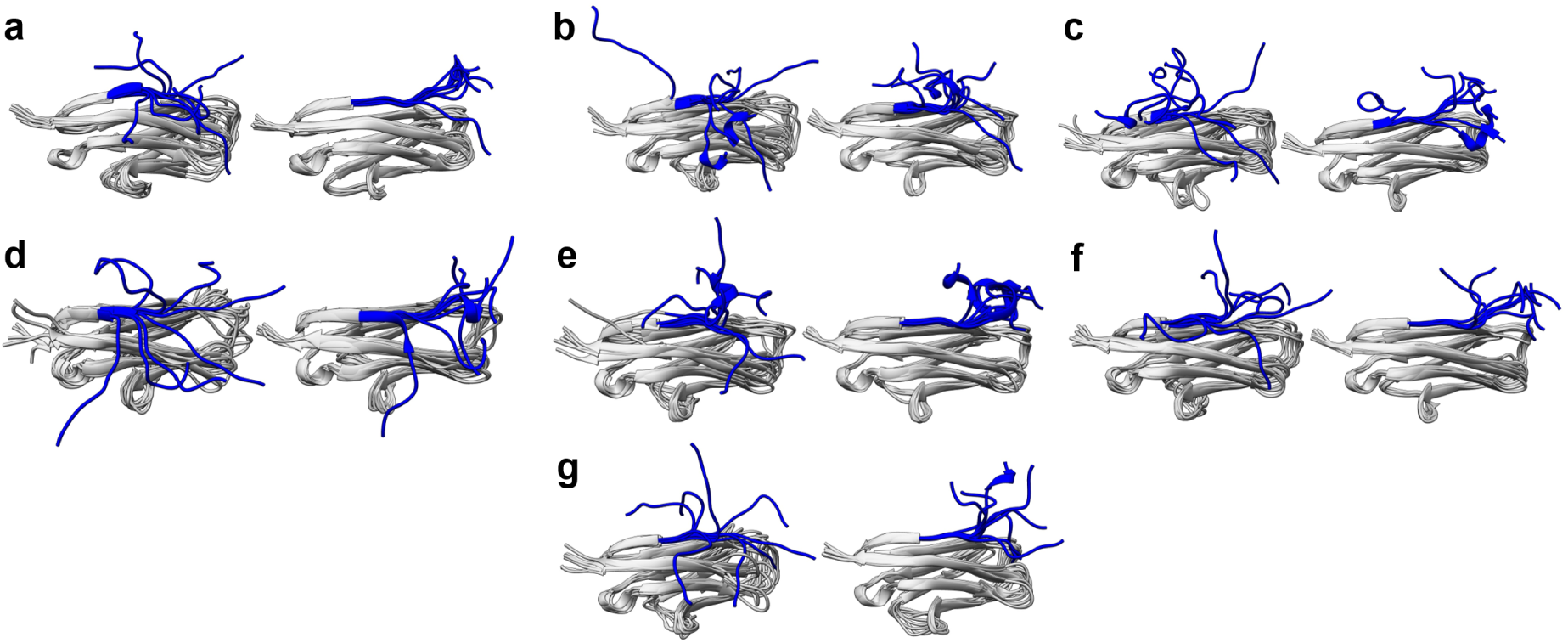
Clusters of the unfolded states of the I91 perturbations for each force field. K-Centers clusters are found to the left and K-Medoids clusters are found to the right. The force fields used are as follows: a) Amber ff99SB-ILDN (in Desmond) b) Amber ff99SB-ILDN (in Amber) c) Amber ff99SB*-ILDN d) Amber FB15 e) Amber ff14SB f) Amber ff14SBonlySC g) Charmm 22*

The observation that the Amber FB15 force field samples a wider space on the Ramachandran map is corroborated by the KCenters algorithm clusters for the Amber FB15 force field (see Figure 4d). The KCenters algorithm for Amber FB15 identifies clusters which exist in a wide variety of different configurations that the other force fields do not seem to sample. This suggests that the Amber FB15 force field does increase the space which the perturbations sample. Overall, it can be seen that most structures sample close to the native structure as seen through the KMedoids algorithm. This suggests that the pathways for the folding of the I91 perturbation are limited. Thus, the I91 perturbation is a useful tool for examining the refolding of peptides because force fields must go through one pathway in order to find the correct native structure.

### Ac-(AAQAA)_3_-NH_2_ Mean Helical Lifetime Can Prove Useful In Identifying Force Field Discrepancies

To examine whether the results of I91 perturbed intermediate refolding could be replicated in other peptides and linked to experimental observables, we also ran Ac-(AAQAA)_3_-NH_2_ equilibration simulations. Previously, the Ac-(AAQAA)_3_-NH_2_ peptide was found to form low amounts of helical structure that sharply decreased as temperature increased. ^32,50^ By running equilibration simulations under previously described conditions ^30,32^ we hoped to to examine how our simulation methods compared against previously published results as well as add data for newly tested force fields. Furthermore, we hoped that examining Ac(AAQAA)_3_-NH_2_ refolding patterns might provide further insight into possible force field discrepancies.

We first analyzed the helical frequency of each residue in the same manner as performed in previous literature.^30,32,32,44^ We found that the Amber ff99SB*-ILDN and Amber FB15 force fields accurately reproduce the 19% helical frequency per residue as observed by NMR experiments (see Figure 5). ^34,50^ Furthermore, the Amber ff99SB-ILDN force fields for both Desmond and Amber do not produce many helices, hovering at less than 10% for each residue. This supports the accuracy of our simulation methods as it mirrors previously published results.^32,34^ The Amber ff14SBonlySC force field also produces this low amount of helical content which is understandable since the dihedral terms were not modified for this force field and should be identical to that of Amber ff99SB-ILDN.

**Figure 5:**
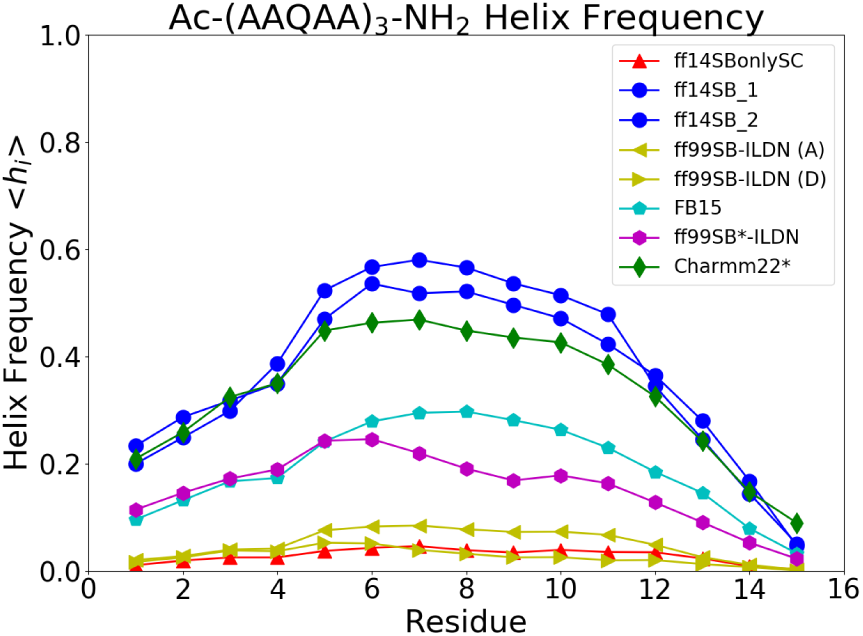
Ac-(AAQAA)_3_-NH_2_ helicity per residue. The helicities were determined with previously established criteria ^32^

In contrast to the low helicities of the Amber ff99SB-ILDN and Amber ff14SBonlySC force fields, the Amber ff14SB force field displays a sharp increase in the amount of helical content. This is corroborated by the I91 perturbed intermediate model which suggested that the empirical dihedral correction for Amber ff14SB created an unexpectedly deep minima in the alpha helical region of the Ramachandran plot. More surprisingly however, the Charmm 22* force field possesses a helical frequency very similar to that of Amber ff14SB. Such a result is interesting because the Charmm 22* and Amber ff14SB force fields display drastically different refolding pathways in the I91 perturbation. Specifically, the Amber ff14SB force field displayed large amounts of helices during I91 perturbation refolding while the Charmm 22* force field did not (see Figure 3). Why then, would the Charmm 22* force field not form helices during I91 perturbation refolding when it does form a large amount of helices in the Ac-(AAQAA)_3_-NH_2_ peptide?

Although different force fields may have similar overall helical frequencies, how those aggregate helical frequencies are created might differ. A force field with two stretches of very stable helix and a force field with lots of small stretches of unstable helix may end up with the same helical frequency over a long period of time. Thus we examined the change in helicity with respect to time to provide a more detailed analysis of the helical frequencies found during Ac-(AAQAA)_3_-NH_2_ equilibration (see Figure 6).

**Figure 6:**
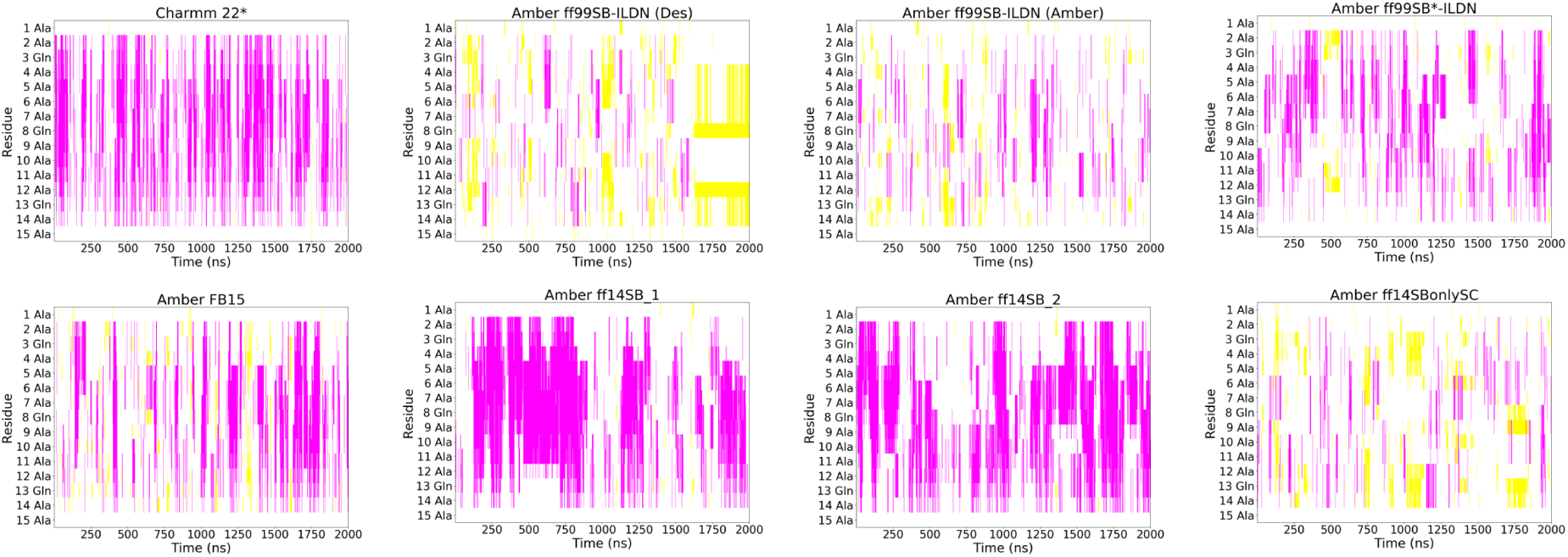
Ac-(AAQAA)_3_-NH_2_ secondary structure formation with respect to time for each force field. Magenta represents helical structures, yellow represents helical structures, and white represents coiled structures. Secondary structures were generated using DSSP with abbreviated criteria^58^

From the graphs of secondary structure with respect to time (see Figure 6) it can be seen that while the Amber ff14SB force field possesses larger amounts of stable helices, the Charmm 22* force field possesses almost as many helices except that they form and break more quickly. This suggests that the primary difference between the Amber ff14SB and Charmm 22* force fields is that the Charmm 22* force field does not stabilize formed helices nearly as well as the Amber ff14SB force field does. It also suggests that the Charmm 22* force field is more likely to initiate the formation of helices than the Amber ff14SB force field.

Additionally, the Amber ff99SB*-ILDN force field also displays large intervals of stable helical content interspersed with smaller amounts of helices. Similar behavior can be found in the Amber FB15 force field (see Figure 6). From these graphs it might be suggested that the lack of helical initiation might be what separates the Amber FB15 and Amber ff99SB*ILDN force fields from the Charmm 22* force field rather than the stability of individual helices.

Another comment must be made on the Amber ff99SB-ILDN force fields for Desmond and Amber. In a large portion of the Desmond simulation, the Ac-(AAQAA)_3_-NH_2_ peptide forms a beta hairpin which does not break. This is not expected since the Ac-(AAQAA)_3_-NH_2_ peptide should form alpha helices. However, we believe that the discrepancy is the result of either sampling or the use of the NMA cap instead of the NH_2_ cap in the peptide simulated using Desmond rather than any differences between the simulation packages. This is because Amber ff99SB-ILDN has already been shown to possess extremely low amounts of helical content. Therefore, it could be suggested that this might translate to an observation of a beta structure. Furthermore, although less likely, the presence of an NMA cap could also create differences in folding behavior.^19^

To quantify differences in helical content over time we counted the helices formed in the secondary structure vs time graphs created using DSSP (see Figure 7). After identifying each helix that was formed, the mean helical lifetime was calculated for each force field. Furthermore, the mean helical lifetime of each residue of Ac-(AAQAA)_3_-NH_2_ was also determined. We performed this extra analysis because one helical region may have individual residues breaking apart and reforming. In such cases, examining the mean helical lifetime of each residue would give a better picture of the dynamics of individual amino acids in a helix for a given force field. The max helical lifetimes overall and for each residue were also counted because the distribution of helix lifetimes is skewed toward very short helical lifetimes and the maximum helical lifetimes gives an idea of the range of helical lifetimes found for each force field.

**Figure 7:**
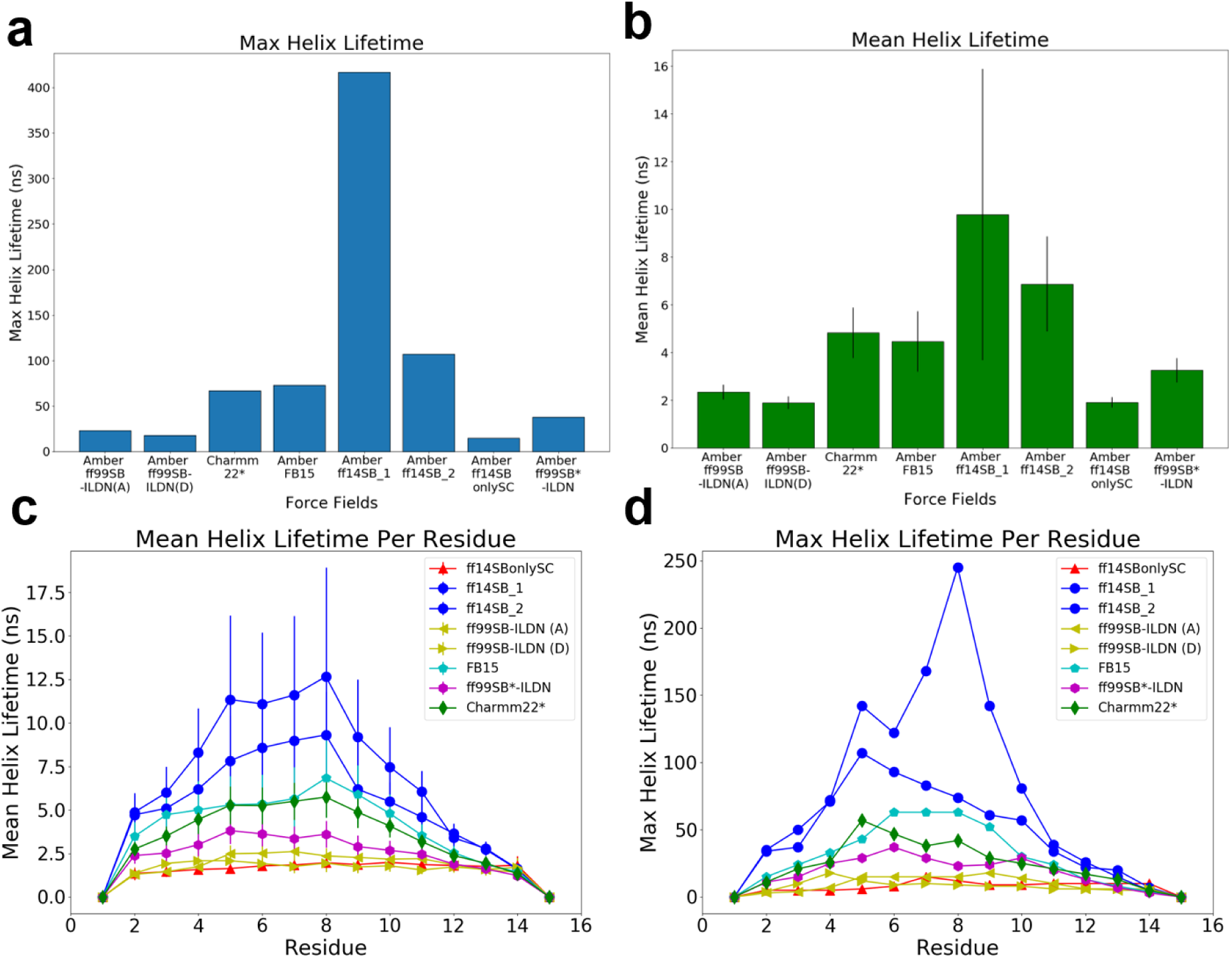
Examination of differences in Ac-(AAQAA)_3_-NH_2_ helical lifetimes. A) The mean helix lifetime for each force field B) The maximum helix lifetime for each force field C) The mean helix lifetime for each residue in each force field D) The maximum helix lifetime for each residue in each force field

Calculating helical lifetimes confirms several observations gained qualitatively from observing the secondary structure of Ac-(AAQAA)_3_-NH_2_ over time (see Figure 6). By examining helical lifetime rather than overall helicity, the helicity of Charmm 22* lowers considerably (see Figure 7). While examining helical frequency, Charmm 22* would have similar frequencies to Amber ff14SB. When examining mean helical frequency however, the Charmm 22* equilibration simulation of Ac-(AAQAA)_3_-NH_2_ reveals helical lifetimes equal to or lower than Amber FB15, a force field which had helical frequency about equal to the experimentally parameterized value. This suggests that Charmm 22* is increasing the initiation of helix formation rather than maintaining helical stability. Although the error bars of the mean helix lifetimes between Charmm 22* and Amber ff14SB overlap, a t-test combining both Amber ff14SB simulations gives a p-value of 0.057 which we believe is likely to be significant.

The difference in mean helical lifetimes does not explain the difference in I91 perturbation refolding of the Amber ff99SB*-ILDN force field compared to the Amber FB15 or Charmm 22* force fields. This can be seen in the analysis of differences in helical lifetimes. While the Charmm 22* and Amber FB15 force fields have similar lifetimes to that of Amber ff99SB*-ILDN, the Amber ff99SB*-ILDN force field forms far more helices than Charmm 22* or Amber FB15 in the I91 perturbed unfolded state. This is likely due to the difference in sequences between the two simulated structures. While the Ac-(AAQAA)_3_-NH_2_ peptide does not contain any lysine or glutamic acid residues, the I91 perturbation contains two glutamic acid residues and one lysine. These residues have been shown to be overly helical in the Amber ff99SB* force field. ^44^ Since the Amber ff99SB*-ILDN force field does not reparameterize the lysine or glutamic acid residues we would expect this result to be transferable. Thus, the change in systems may explain why helical constructs are observed in the I91 perturbation but not during Ac-(AAQAA)_3_-NH_2_ equilibration simulations.

To examine force field parameterization differences which might lead to differing Ac-(AAQAA)_3_-NH_2_ and I91 perturbation folding results, we constructed potentials of mean force for each of the Ac-(AAQAA)_3_-NH_2_ simulations. Through these potential of mean force plots we hoped to identify regions of the Ramachandran plot which a force field might sample that might explain the discrepancies between the I91 Perturbation and Ac-(AAQAA)_3_-NH_2_ results (see Figure 8).

**Figure 8:**
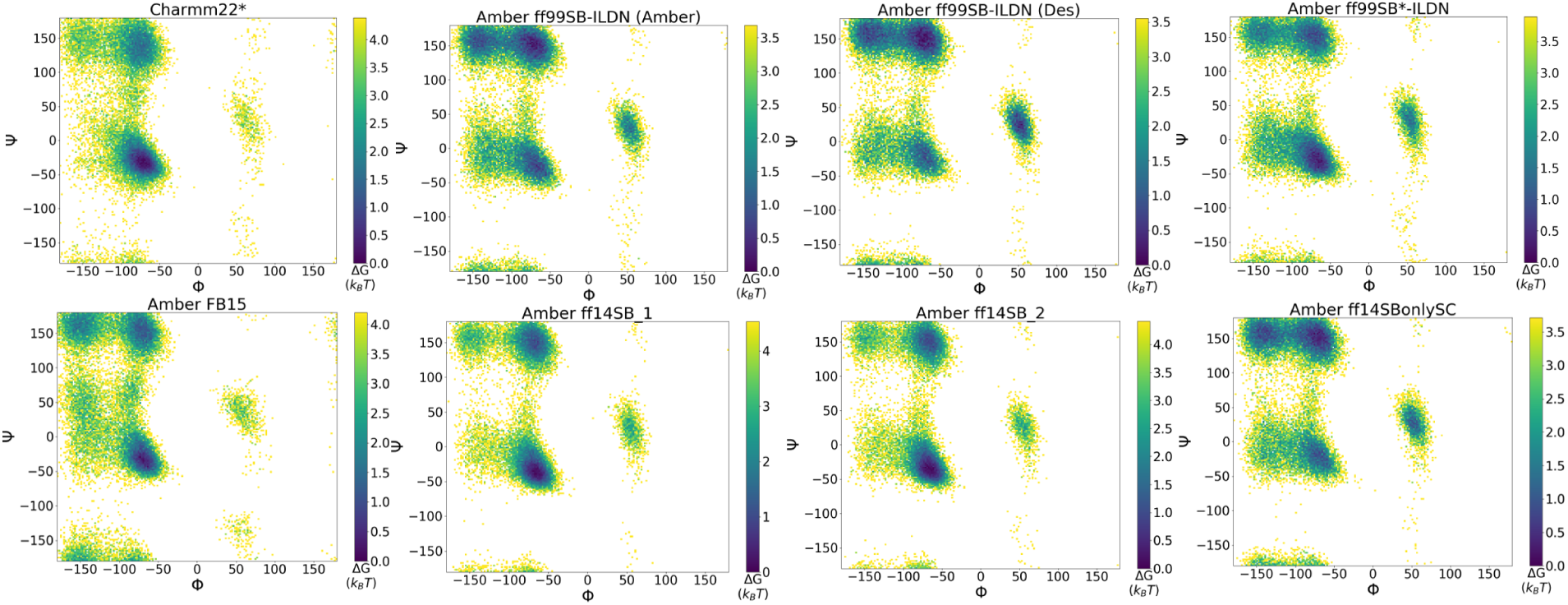
Ac-(AAQAA)_3_-NH_2_ potentials of mean force created from the dihedral angles found in equilibration simulations in each of the observed force fields

One notable difference occurs between the helical region of the Charmm 22* force field and the Amber force fields. While the Charmm 22* force field possesses one minimum for the alpha helical region of the Ramachandran plot, all Amber force fields possess at least two minima for the alpha helical region. All of the force fields tested have a minimum at the a_*h*_ region: *φ ϵ*[−100°, −30°] and *ψ ϵ*[−67°, −7°]. Amber force fields however, possess an additional region of low energy found between *φ ϵ*[-160°,-100°] and *ψ ϵ*[−50°, 50°] which is around 2 *k*_*B*_T greater than the minimum found in the a_*h*_ region (see Figure 2 and Figure 8).

Additionally, in the area between the beta and alpha regions (*ψ ϵ*[25°,75°]), the Charmm 22* force field potential of mean force plot possesses a gradient that is much more widespread and evenly distributed than in any of the other potential of mean force plots (see Figure 8). We further note that the area between the alpha and beta region varies between each of the Amber force fields. The Amber FB15 force field appears to have the most widespread sampling of the dihedral angles in this region while the Amber ff14SB force field samples the region the least. Indeed, the Amber ff14SB force field barely samples the region *φ ϵ* [-160°,-100°] and *ψ ϵ*[−50°, 50°] likely as a result of the increased barrier provided by the Amber ff14SB empirical backbone dihedral correction. ^33^

The differences in sampling in the area between the beta and alpha region and in the extra minimum found in Amber forcefields suggests a possible mechanism for the discrepancy in helix formation for the Charmm 22* force field. Whereas other force fields possessed much greater barriers, the lack of phi barriers to the alpha region of the Charmm 22* force field allows for an easy transition between beta and alpha regions thereby increasing the rate of helix formation by the Charmm 22* force field. In contrast, the Amber ff14SB force field possesses nearly no points in half of the transition region (see Figure 8), illustrating the difficulties of moving from the alpha region to the beta region in the Amber ff14SB force field as compared to the Charmm 22* force field.

## Discussion and Conclusion

In this paper we examined how various force fields could affect the refolding of the I91 mechanically perturbed intermediate and the helical peptide Ac-(AAQAA)_3_-NH_2_. By examining the distribution of dihedral angles, the frequency of secondary structures, and the arrangement of clustered structures we observed the increased formation of helices in the unfolded state for the Amber ff14SB and Amber ff99SB*-ILDN force fields. Additionally we observed that even though the Charmm 22* force field had similar overall helical frequencies to the Amber ff14SB force field, that the Charmm 22* force field possesses a much smaller mean helical lifetime.

The Amber ff99SB*-ILDN and Amber ff14SB force fields both displayed difficulty refolding the I91 mechanically perturbed intermediate. The difference is particularly interesting since these force fields form helices in the unfolded state of the I91 perturbation (see Figure 3). In the native state however, the I91 protein perturbation forms a beta strand and thus forms native contacts with other beta strands. In contrast, the presence of helical intermediates for the small perturbation results in the creation of a significant amount of non-native contacts. Since the I91 perturbation is known to refold experimentally, ^38^ and given the idea of downhill folding, ^29,43^ it is unlikely that the helical constructs of the unfolded state contribute to the folding pathway and rather represent a local minima which the perturbation gets trapped in.

Although the Amber ff99SB*-ILDN and Amber ff14SB force fields both display difficulties refolding the I91 perturbation, the reasons for these difficulties likely differ. It has been shown that the Amber ff99SB* force field is overly helical for glutamic acid and lysine residues. ^44^ This can be seen in how both the Amber ff14SBonlySC and Amber FB15 force fields, which do reparameterize the glutamic acid and lysine side chain torsion angles, can more robustly refold the I91 perturbation than the Amber ff99SB*-ILDN force field (see Table 1).

The Amber ff14SB force field also displays difficulties refolding the I91 perturbation. However, the force field displays higher amounts of helical content than all of the other tested Amber force fields. Furthermore, the Amber ff14SB force field likely does not contain incorrect helicities for glutamic acid and lysine as it is based on the Amber ff14SBonlySC force field which reparameterized all side chains and robustly refolds the I91 perturbation (see Table 1). Thus, the only difference that could cause much greater helicites for the Amber ff14SB force field is the Amber ff14SB empirical backbone dihedral correction. Although this correction has been validated for other small peptides, ^33^ the failure of this correction to match experimentally observed helicities in the Ac-(AAQAA)_3_-NH_2_ peptide as well as its difficulty refolding the I91 perturbation suggest that the correction overstabilizes helices. These observations have been replicated in implicit solvent,^14^ although it should be noted that such observations may not be reliable as the dihedral correction for ff14SB was intended for explicit solvent simulations. ^33^ Furthermore, such an increase in helicities was not replicated in an analysis of A*β*_16*−*22_ dimer. We note that these discrepancies may be explained however, by the aforementioned study examining dimerization instead of refolding and also the more limited time scales per simulation (200 ns vs 2 *µs*).^25^

The ease of Amber ff14SBonlySC and Amber FB15 in refolding the I91 perturbation also raises potential questions about how parameterization affects protein behavior. From the dihedral maps of the I91 mechanical perturbation unfolding state it can be observed that there are noticeable gaps in the beta and alpha region for the Amber ff14SBonlySC force field while the AMBER FB15 force field samples the entire region very widely. This presents two different mechanisms for the efficacy of these force fields in refolding the I91 protein perturbation. The Amber ff14SBonlySC potential of mean force plots along with its low helical frequency during Ac-(AAQAA)_3_-NH_2_ equilibration simulations suggest that the force field is biased toward the beta strand region of the Ramachandran plot. This and the lack of the additional minima in the region suggest that the ff14SBonlySC force field easily forms beta stranded proteins. In contrast, the wide sampling of the Amber FB15 force field suggests that the Amber FB15 force field does not get trapped in any single minima. Instead, the Amber FB15 force field will eventually make its way to the true minima after broadly sampling a larger configuration space.

Furthermore, the difference in helices present in the unfolded state of the I91 perturbation unfolded state suggests that even if two force fields have similar overall helicities as observed with the Ac-(AAQAA)_3_-NH_2_ peptide, the mean helical lifetime can affect whether those helices are actually observed during the refolding pathway. This can be seen when comparing Charmm 22* to Amber ff14SB. Although Charmm 22* Ac-(AAQAA)_3_-NH_2_ equilibration simulations resulted in the same overall helical frequencies as the Amber ff14SB equilibration simulations, the mean helical lifetimes differed. The difference was noticeable in the unfolded state of the I91 perturbation where the Charmm 22* force field did not form many helices while the Amber ff14SB force field did. These observations can be correlated with observations of the potential of mean force plots between the Charmm 22* and Amber force fields. The singular alpha region of the Charmm 22* force field, without the extra minima found in the region (*φ ϵ*[-160°,-100°] and *ψ ϵ*[−50°, 50°]) may result in a more direct transition between the beta and alpha regions of the Ramachandran plot which may result in more frequent but less stable helices as compared to other Amber force fields with similar overall helical frequencies.

The extra data from the I91 perturbed intermediate allowed for the controlled examination of beta strand refolding. The differences in this refolding pathway then led to observations about force field discrepancies that might have been otherwise unnoticed by examining bulk measurements like Ac-(AAQAA)_3_-NH_2_ helical frequency. Thus, investigating force field accuracy through protein structure perturbations provides a way of testing the transferability of parameterization schemes and actual protein folding simulations while still keeping a system small enough to reduce the time and simulation costs of folding. Thus the creation of mechanically perturbed intermediates can prove to be a useful tool in identifying various force field discrepancies.

## Acknowledgement

The authors thank the following funding sources: NSF Grant MCB-1517245 and NSF Grant MCB-1817556, Sigma Xi Grant G2018031596204454, Duke Deans’ Summer Research Fellowship, and NCICU Undergraduate Research Fellowship.

We thank R.B. Best, E. Josephs, and J. Pajak for helpful discussions regarding the nature of this publication. We thank J. Wu for help with the creation and set up of our computational equipment.

Molecular graphics and analyses performed with UCSF Chimera, developed by the Resource for Biocomputing, Visualization, and Informatics at the University of California, San Francisco, with support from NIH P41-GM103311.

## Graphical TOC Entry

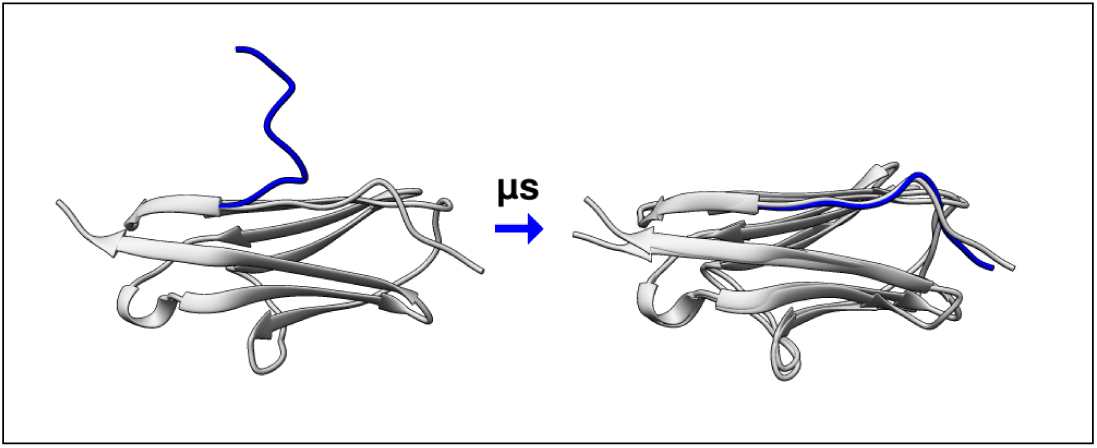

