## Supplemental Information for "Exploiting a Mechanical Perturbation of Titin Domain to Identify How Force Field Parameterization Affects Protein Refolding Pathways"

**Supporting Material**

**Machine Simulation Performance Data**

All simulations were performed using consumer grade hardware and GPU cards.

Timings of simulations are recorded for various GPU cards and CPUs to help gain a sense of what’s currently feasible for most labs when running molecular dynamics simulations without supercomputers. We’ve found that timings can vary by around 10 ns/day per microsecond run.

Simulations using Desmond GPU:

| **System** | **Simulation Details** | **System Size (atoms)** | **GPU** | **CPU** | **Timing (ns/day)** |
| --- | --- | --- | --- | --- | --- |
| I91 Perturbation | Cutoff: 12 Å | 18419 | Nvidia GTX 980 Ti | 1 core, AMD Phenom II 1090T, 3.2 GHz | 205.955 |
| I91 Perturbation | Cutoff: 9 Å | 18419 | Nvidia GTX 980 Ti | 1 core, AMD Phenom II 1090T, 3.2 GHz | 299.032 |
| (AAQAA)_3_ | Cutoff: 9.5 Å | 31485 | Nvidia GTX 980 Ti | 1 core, AMD FX-8350, 4.0 GHz | 204.008 |
| (AAQAA)_3_ | Cutoff: 12 Å | 31485 | Nvidia GTX 980 Ti | 1 core, AMD Phenom II 1090T, 3.2 GHz | 143.686 |

Simulations using Amber GPU:

| **System** | **Simulation Details** | **System Size (atoms)** | **GPU** | **CPU** | **Timing (ns/day)** |
| --- | --- | --- | --- | --- | --- |
| I91 Perturbation | Cutoff: 9.5 Å | 19091 | Nvidia Titan V | 1 core, Intel i5-7600K, 4.098 GHz | 455.26 |
| I91 Perturbation | Cutoff: 9.5 Å | 19091 | Nvidia GeForce 1080 (boosted) | 1 core, Intel i5-7600K, 4.098 GHz | 320.88 |
| I91 Perturbation | Cutoff: 9.5 Å | 19091 | Nvidia GeForce 1080 | 1 core, AMD Phenom II 980, 3.7 GHz | 259.09 |
| I91 Perturbation | Cutoff: 9.5 Å | 19091 | Nvidia GTX 980 Ti | 1 core, AMD FX-8350, 4.0 GHz | 208.60 |
| I91 Perturbation | Cutoff: 9.5 Å | 19091 | Nvidia GTX 980 Ti | 1 core, AMD Phenom II 1090T, 3.2 GHz | 203.37 |
| (AAQAA)_3_ | Cutoff: 9.5 Å | 32367 | Nvidia Titan V | 1 core, Intel i5-7600K, 4.098 GHz | 318.67 |
| (AAQAA)_3_ | Cutoff: 9.5 Å | 31986 | Nvidia GeForce 1080 (boosted) | 1 core, Intel i5-7600K, 4.098 GHz | 201.02 |
| (AAQAA)_3_ | Cutoff: 9.5 Å | 32367 | Nvidia GeForce 1080 (boosted) | 1 core, Intel i5-7600K, 4.098 GHz | 198.58 |

**I91 Perturbation Refolding Data**

The fractional Q-value of a protein was constructed according to the following equation (Best et al., 2013):

$$Q\left( X \right)= \frac{1}{N}\sum_{(i,j)} \frac{1}{1+exp[\beta(r_{ij}\left( X \right)-\lambda r_{ij}^{0})]}$$

All pairs of heavy atoms i and j where one heavy atom must belong to a residue in the perturbation (residues 1-10) were included in the summation. $r_{ij}^{0}$ was taken to be the distance between i and j in the native state (crystal structure 1waa (Stacklies et al., 2009)). $\beta$ is the smoothing parameter which was set as 5 Å^-1^ (Best et al., 2013). $\lambda$ accounts for fluctuations in the contact and was set as 1.8 (Best et al., 2013).

The following Q-value plots were produced and used to identify the unfolded state of the I91 perturbation. The unfolded state was defined as the last point in time where the Q-value was < 0.4. The value of Q < 0.4 was made to prevent inclusion of the transition state into analysis of the unfolded state. A perturbation was considered “folded” if the Q-value exceeded 0.85. The second trial of the ff14SB force field was extended to 5 μs to see whether the force field could eventually refold the perturbation. This was not included in the table in the main text as every trial in the table only encompasses the time up to 2 μs.

Charmm 22* (Trials 1-5):

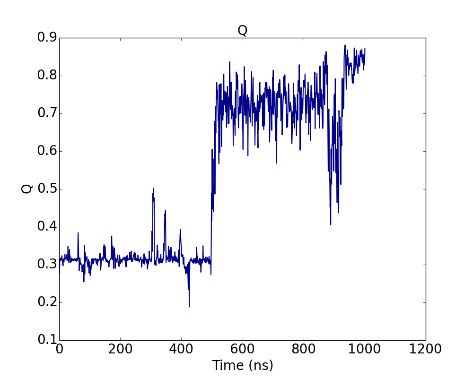

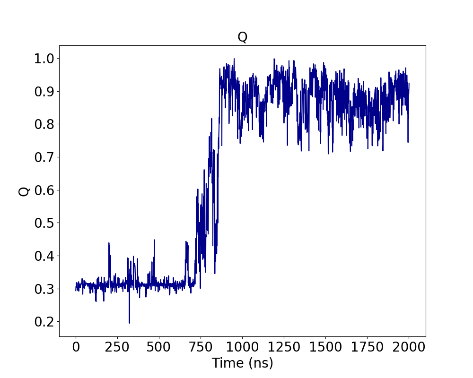

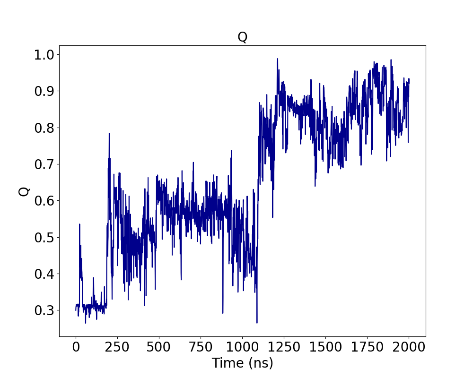

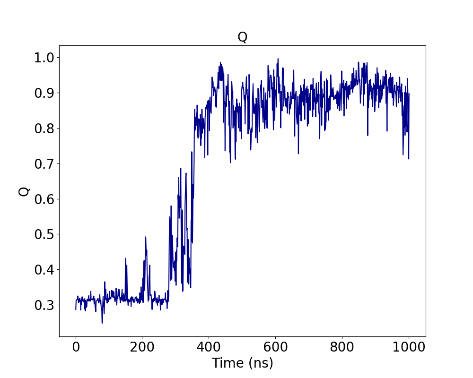

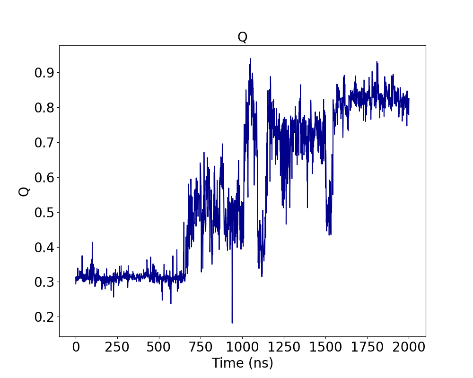

Amber ff99SB-ILDN, Desmond simulated (Trials 1-5):

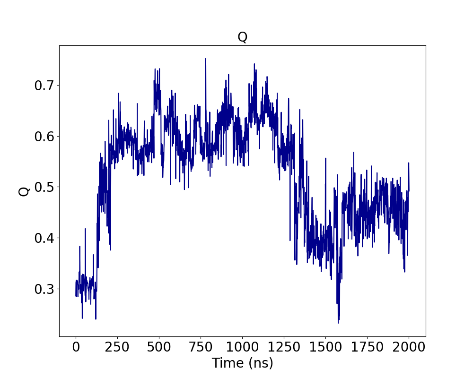

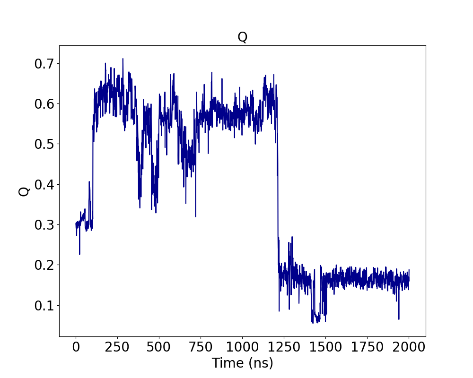

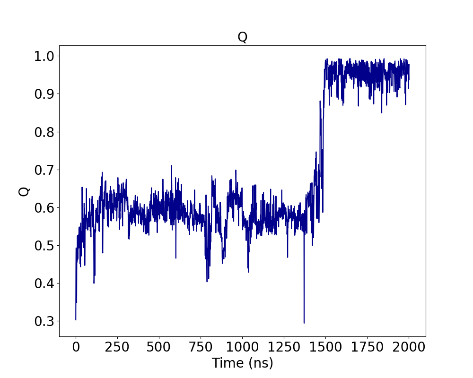

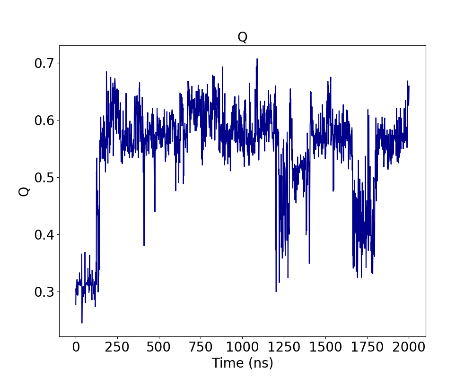

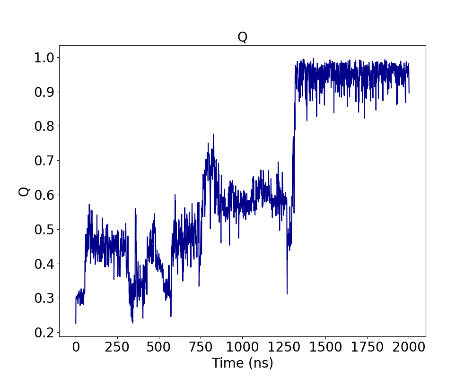

Amber ff99SB-ILDN, Amber simulated (Trials 1-5):

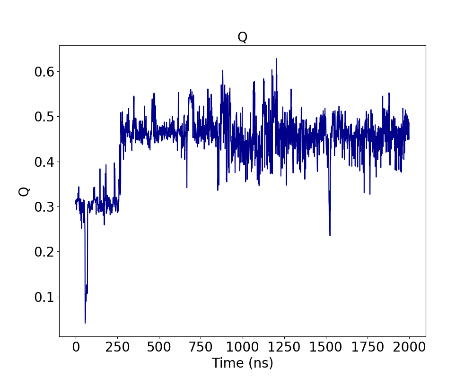

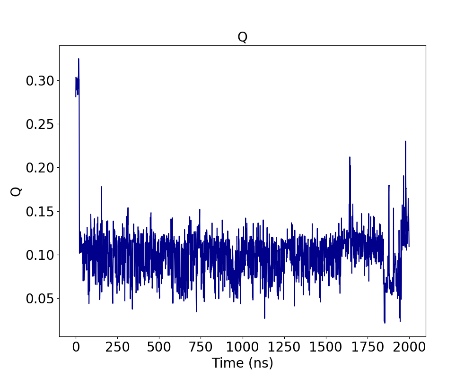

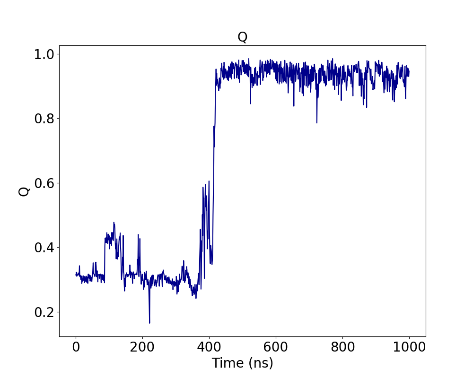

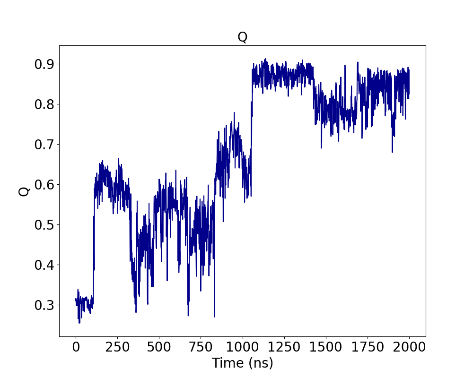

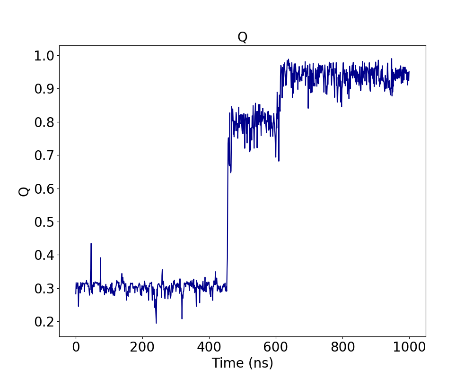

Amber FB15 (Trials 1-5):

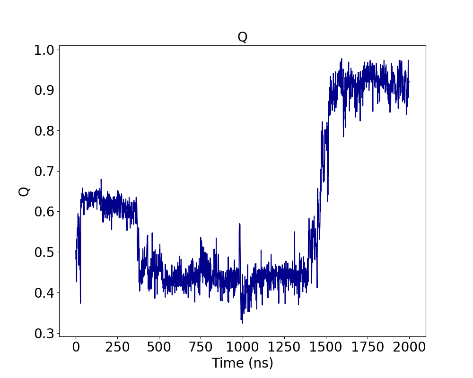

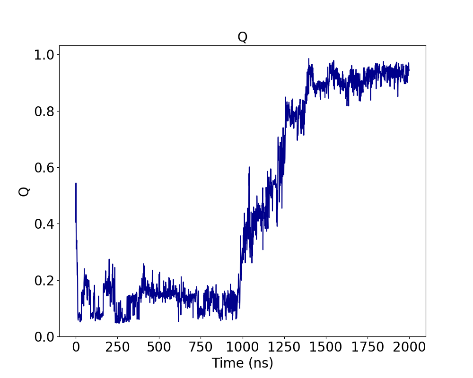

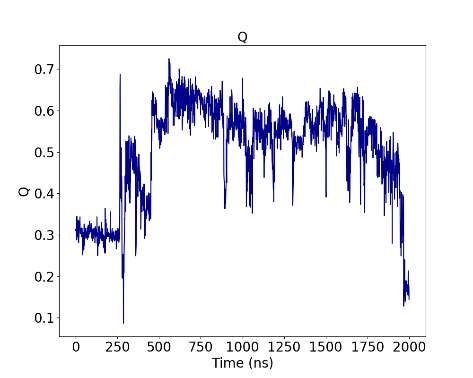

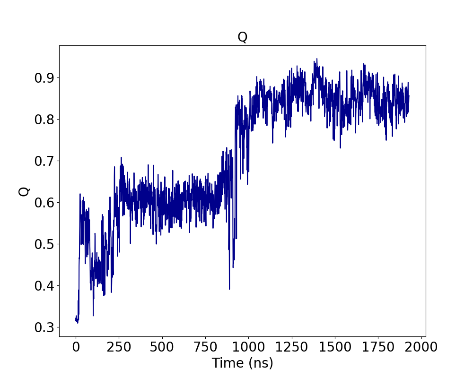

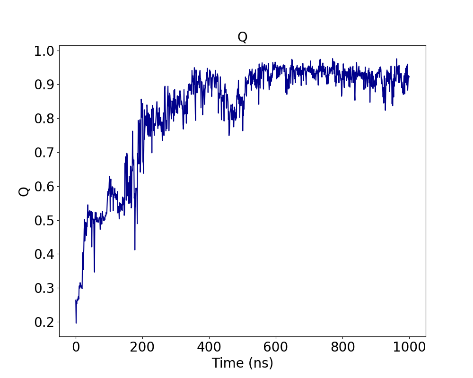

Amber ff14SB:

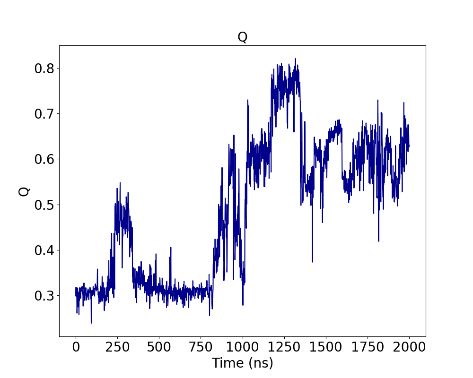

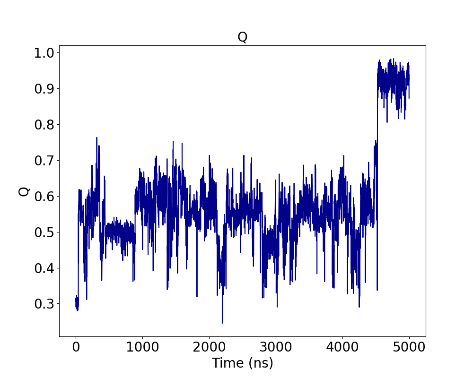

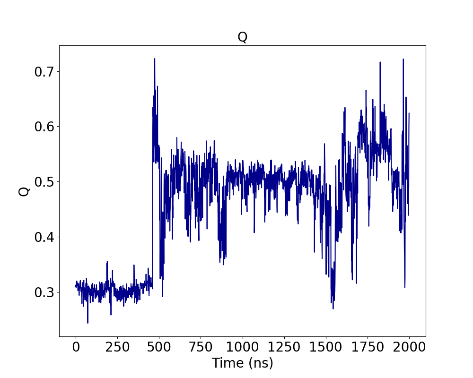

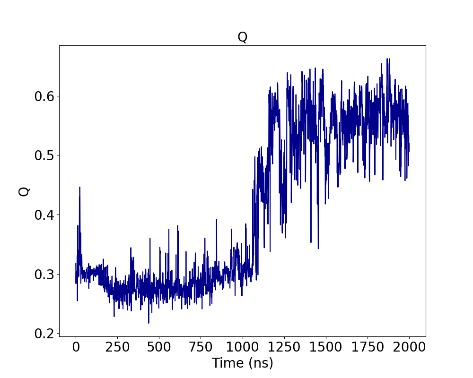

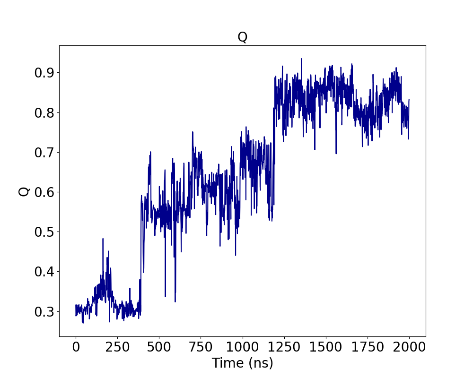

Amber ff14SB without dihedral correction (Trials 1-5):

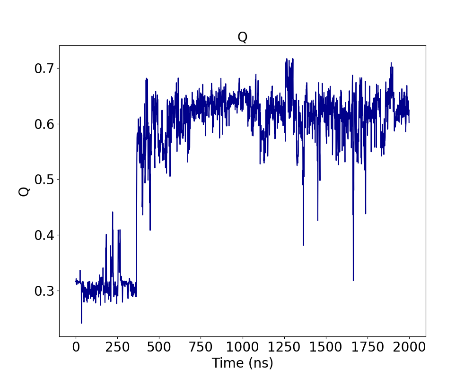

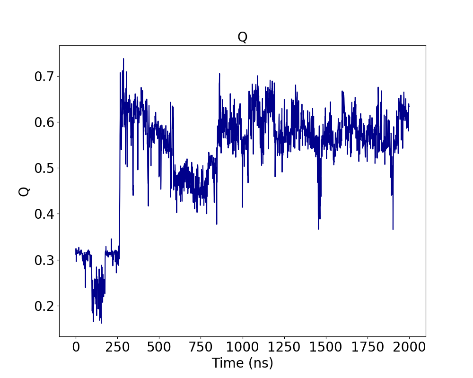

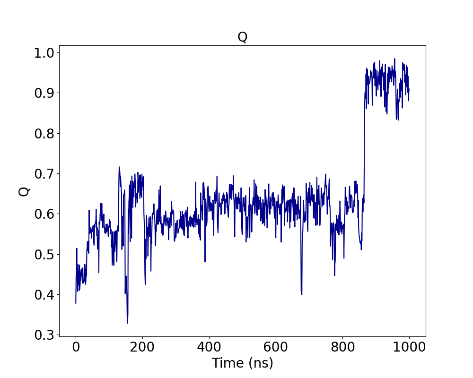

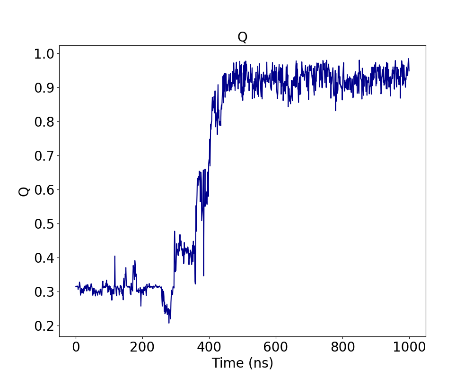

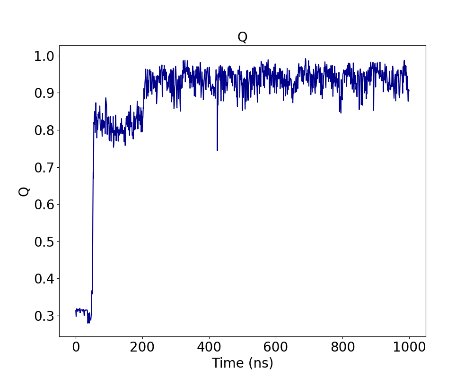

Amber ff99SB*-ILDN (Trials 1-5):

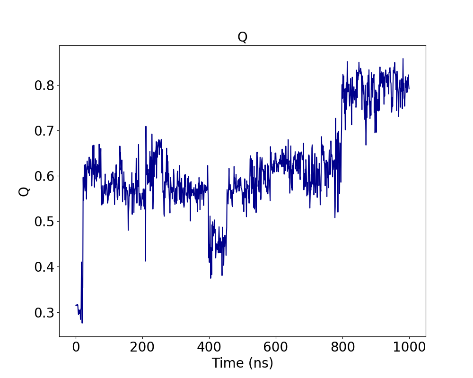

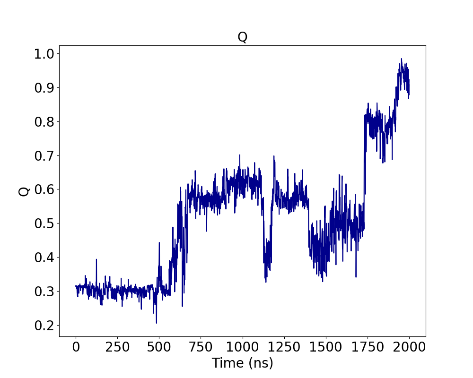

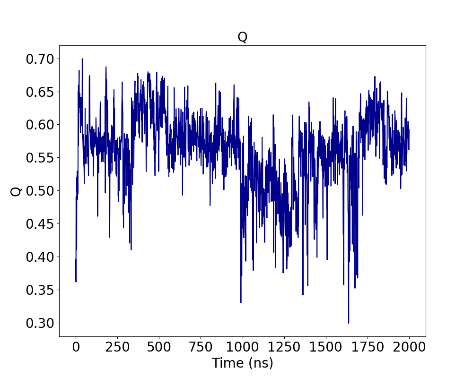

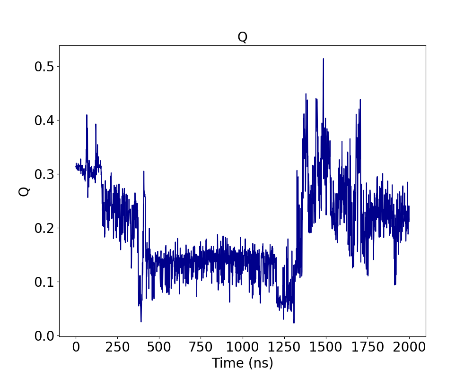

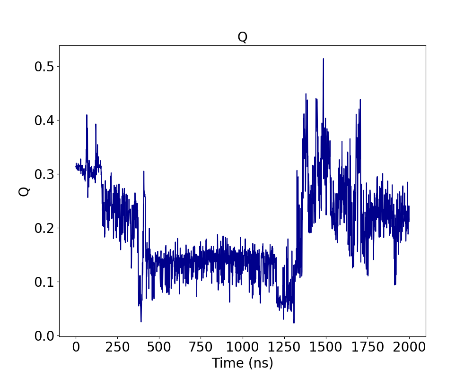

**I91 Refolded Trajectories**

The following figure visualizes the folded state of each trajectory that successfully folded the I91 perturbation. The last frame of each trajectory that successfully folded the I91 perturbation is taken (colored in blue) and aligned with the crystal structure of the native state (colored in gray).

The force fields are arranged as following: a. Amber ff99SB-ILDN, Desmond simulated b. Amber ff99SB-ILDN, Amber simulated c. Amber ff99SB*-ILDN d. Amber FB15 e. Amber ff14SB f. Amber ff14SB without dihedral correction g. Charmm 22*

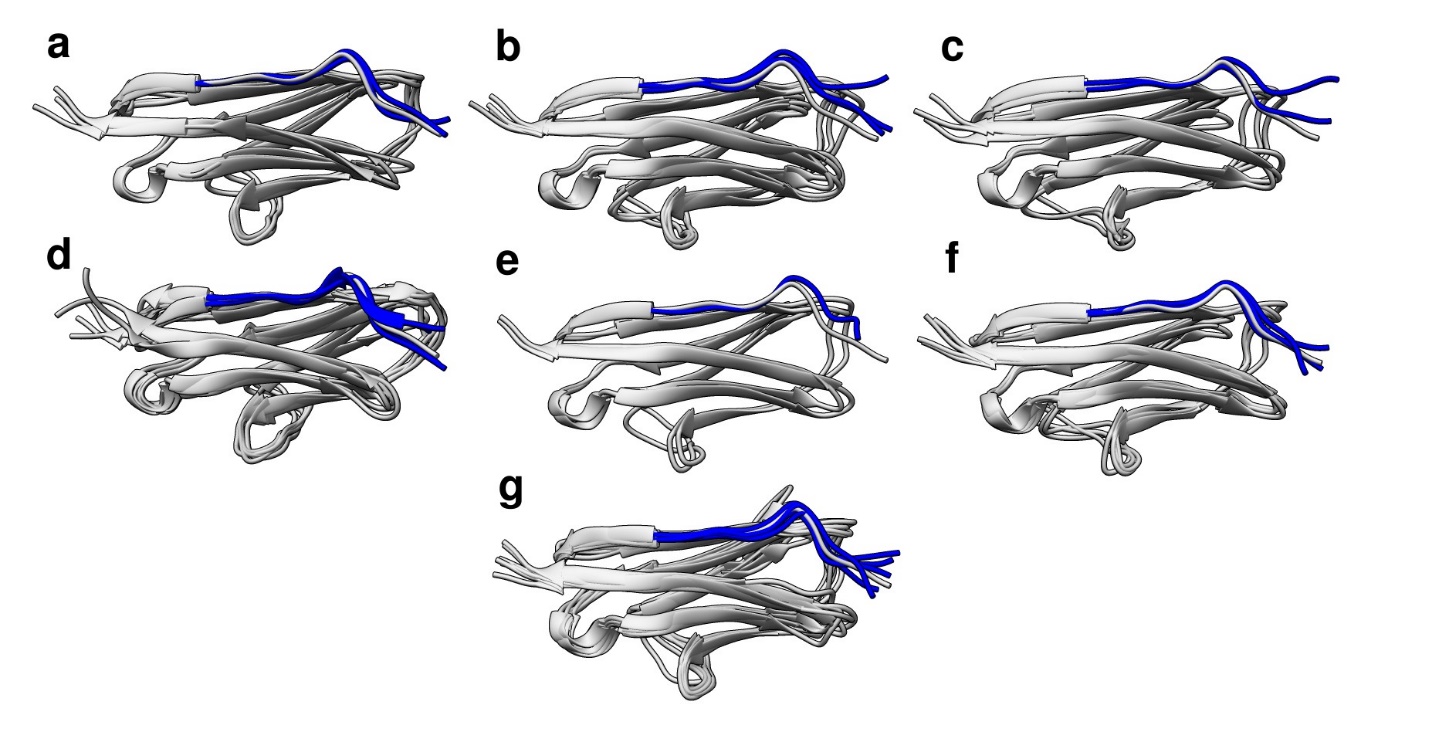

**Analysis of Helix Lifetimes in Ac-(AAQAA)_3_-NH_2_:**

We defined the lifetime of a helix as the length of time which a helical residue or its adjacent counterparts lasted. The breakpoint was defined as the point in time where all residues found in the helix were no longer helical. An example of a breakpoint is highlighted by the red circle:

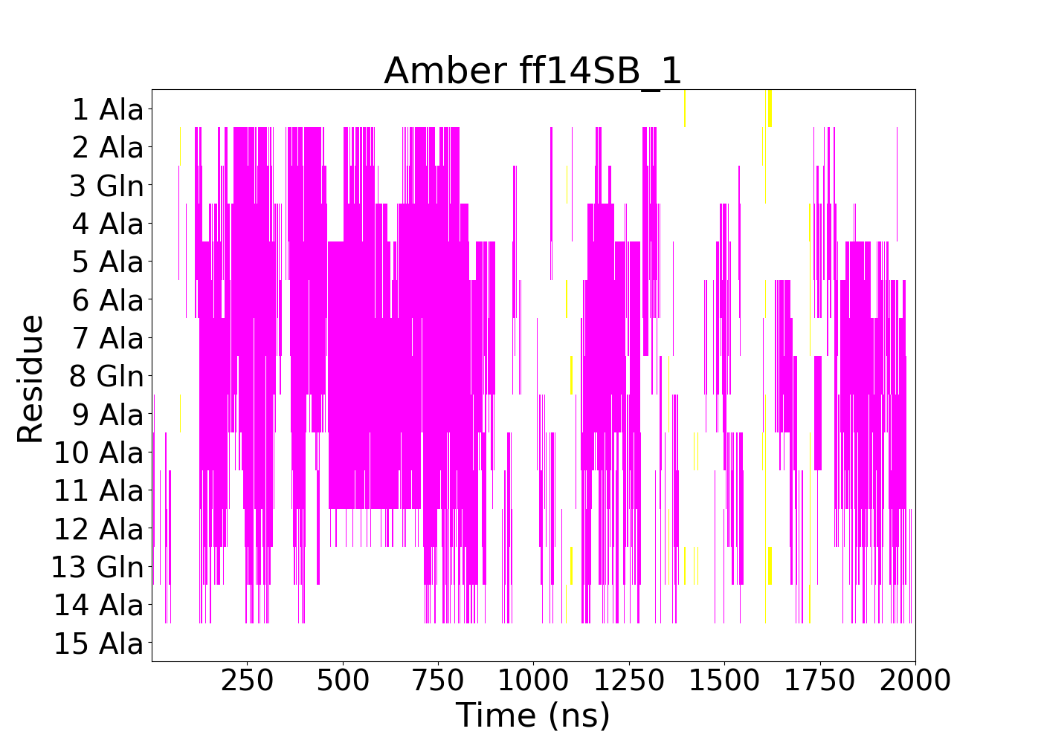

In order to identify helices using this definition we performed a breadth first search on the matrix created by the residue configurations over time. Every unique breadth first search that could be conducted was considered a “helix”. The code for a search using the DSSP implementation of mdtraj (McGibbon et al., 2015) is given as follows:

\# Import statements

import mdtraj as md

import numpy as np

import matplotlib as mpl

from matplotlib import pyplot as plt

from matplotlib import colors

\# Load trajectory and topology

traj = md.load('desmond_md_job_1/desmond_md_job_1_trj/clickme.dtr',top = 'desmond_md_job_1/desmond_md_job_1-out.pdb')

prot = traj.atom_slice(traj.topology.select("protein"))

\# Compute DSSP criteria

dssp = md.compute_dssp(prot)

[t,z] = dssp.shape # get time frame of data

protDSSP = dssp[:,1:16].copy() # get slice of DSSP matrix involving actual protein

\# create copy of numpy

searched = np.zeros(protDSSP.shape)

\# create a list of helices and a list of interval lengths

helices = []

intervals = []

\# search through the matrix of helices and times, iterate by row first

for i in range(0,t):

for resid in range(0,15):

if (protDSSP[i,resid] == 'H') and (searched[i,resid] == 0):

\# create helix

helix = np.zeros(protDSSP.shape)

\# counter for adding to residues

begin = i

end = i

index = (i,resid)

queue = []

queue.append(index)

\# update individual helix and searched matrix

helix[index] = 1.0

searched[index] += 1.0

while queue:

\# Dequeue a vertex from queue and print it

s = queue.pop(0)

searched[s] += 1.0

\# Check whether the end and beginning interval can be changed

if (s[0] > end):

end = s[0]

if (s[0] < begin):

begin = s[0]

if (s[0] == 0) and (s[1] == 0):

if (protDSSP[s[0]+1,s[1]] == 'H') and (helix[s[0]+1,s[1]] == 0):

newIndex = (s[0]+1,s[1])

queue.append(newIndex)

helix[newIndex] += 1.0

if (protDSSP[s[0],s[1]+1] == 'H') and (helix[s[0],s[1]+1] == 0):

newIndex = (s[0],s[1]+1)

queue.append(newIndex)

helix[newIndex] += 1.0

elif (s[0] == 0) and (s[1] == 14):

if (protDSSP[s[0]+1,s[1]] == 'H') and (helix[s[0]+1,s[1]] == 0):

newIndex = (s[0]+1,s[1])

queue.append(newIndex)

helix[newIndex] += 1.0

if (protDSSP[s[0],s[1]-1] == 'H') and (helix[s[0],s[1]-1] == 0):

newIndex = (s[0],s[1]-1)

queue.append(newIndex)

helix[newIndex] += 1.0

elif (s[0] == t-1) and (s[1] == 0):

if (protDSSP[s[0],s[1]+1] == 'H') and (helix[s[0],s[1]+1] == 0):

newIndex = (s[0],s[1]+1)

queue.append(newIndex)

helix[newIndex] += 1.0

elif (s[0] == t-1):

if (protDSSP[s[0]-1,s[1]] == 'H') and (helix[s[0]-1,s[1]] == 0):

newIndex = (s[0]-1,s[1])

queue.append(newIndex)

helix[newIndex] += 1.0

if (protDSSP[s[0],s[1]+1] == 'H') and (helix[s[0],s[1]+1] == 0):

newIndex = (s[0],s[1]+1)

queue.append(newIndex)

helix[newIndex] += 1.0

if (protDSSP[s[0],s[1]-1] == 'H') and (helix[s[0],s[1]-1] == 0):

newIndex = (s[0],s[1]-1)

queue.append(newIndex)

helix[newIndex] += 1.0

elif (s[0] == 0):

if (protDSSP[s[0]+1,s[1]] == 'H') and (helix[s[0]+1,s[1]] == 0):

newIndex = (s[0]+1,s[1])

queue.append(newIndex)

helix[newIndex] += 1.0

if (protDSSP[s[0],s[1]+1] == 'H') and (helix[s[0],s[1]+1] == 0):

newIndex = (s[0],s[1]+1)

queue.append(newIndex)

helix[newIndex] += 1.0

if (protDSSP[s[0],s[1]-1] == 'H') and (helix[s[0],s[1]-1] == 0):

newIndex = (s[0],s[1]-1)

queue.append(newIndex)

helix[newIndex] += 1.0

else:

if (protDSSP[s[0]-1,s[1]] == 'H') and (helix[s[0]-1,s[1]] == 0):

newIndex = (s[0]-1,s[1])

queue.append(newIndex)

helix[newIndex] += 1.0

if (protDSSP[s[0],s[1]-1] == 'H') and (helix[s[0],s[1]-1] == 0):

newIndex = (s[0],s[1]-1)

queue.append(newIndex)

helix[newIndex] += 1.0

if (protDSSP[s[0]+1,s[1]] == 'H') and (helix[s[0]+1,s[1]] == 0):

newIndex = (s[0]+1,s[1])

queue.append(newIndex)

helix[newIndex] += 1.0

if (protDSSP[s[0],s[1]+1] == 'H') and (helix[s[0],s[1]+1] == 0):

newIndex = (s[0],s[1]+1)

queue.append(newIndex)

helix[newIndex] += 1.0

interval = (begin,end)

intervals.append(interval)

helices.append(helix)

elif(searched[i,resid] == 0):

searched[i,resid] += 1.0

**Achieving -NH_2_ capping using Charmm 22* and further analysis:**

An elongated structure of the Ac-(AAQAA)_3_-NH_2_ peptide was created using tleap and saved as a pdb file. Inside the pdb file, the -NH_2_ cap is initially labeled “NHE” with an amino acid number of 17.

Although the Charmm 22* force field does not have an explicit -NH_2_ cap, it does have a template for Alanine with a terminal amide group. This template is labeled “ALA_CT2”, and contains the following atoms:

["NT", 7, -0.62, ["NH2"]],

["HT1", 1, 0.32, ["H"]],

["HT2", 1, 0.30, ["H"]]

In order to match the ALA_CT2 template found in the Charmm 22* force field, we changed the name of the “NHE” residues to “ALA”, renumbered the residue number from 17 to 16, and relabeled the atoms as “NT”, “HT1”, and “HT2” respectively. A confirmation message produced during Viparr: “Note: Struct 2, Res <" ",16> (ALA) matched (ALA_CT2)” gives us confidence that the template was successfully matched.

In order to conduct dihedral analysis using VMD, we changed the last three residues back to “ALA” with residue number 17 instead of 16.
